# Identification of five characteristic ferroptosis-related genes as novel biomarkers for ischemic stroke using bioinformatic and machine learning strategies

**DOI:** 10.1101/2023.07.20.549866

**Authors:** Xin Feng, Chunhua Huang, Lin Ye, Zhongbo Xu

## Abstract

Ferroptosis is crucial in neuronal cell death, associated with various neurological disorders. However, the role of ferroptosis-related genes (FRGs) in ischaemic stroke (IS) has yet to be well elucidated. We downloaded IS-related gene information and FRGs from the Gene Expression Omnibus (GEO) and Ferritin databases (FerrDb). 22 IS-related DE-FRGs were obtained. Functional enrichment analysis showed that these genes involve multiple regulatory pathways related to IS pathogenesis, including redox, amino acid metabolism, and cell cycle. Subsequently, DUOX2, MDM2, EGFR, MAP3K14, and TRIM46 were identified as core marker genes among the 22 DE-FRGs by lasso and SVM-RFE algorithms. The construction of the nomogram using the five marker genes had excellent diagnostic value. In addition, CIBERSORT analysis showed that changes in the immune microenvironment of IS patients might be associated with TRIM46, MDM2, and DUOX2. In addition, a total of 66 drugs targeting two characteristic FRGs were obtained. The ceRNA networks revealed complex regulatory relationships based on characteristic FRGs. These findings provide new insights into the diagnosis and treatment of IS.

## Introduction

Ischemic stroke (IS) is a common cerebrovascular disease caused by the blockage of blood flow to the neck or brain[1]. It often develops acutely, has a narrow therapeutic window, and poses a significant threat to human health. Studies have shown that since 2010, stroke has become the second leading cause of death worldwide after tumors, with a mortality rate of over 11% and an increasing trend with age[2]. The severe cognitive and motor impairment caused by ischaemic stroke significantly burdens families and society[3]. The rapid onset of the disease and the many delays in treatment make early thrombolytic therapy challenging to administer, thus affecting the outcome[4]. In addition, even if revascularisation is completed within the treatment window, it may lead to more serious adverse effects, such as conversion to cerebral hemorrhage [5]. Therefore, exploring the pathogenesis of ischaemic stroke and new biomarkers and therapeutic targets is essential.

Ischemic stroke is caused by a localized lack of blood supply to the brain tissue, secondary to neuronal cell death, a process in which multiple cell death pathways are involved[6]. Cell death pathways are complex, and the common form of programmed cell death is apoptosis, in addition to other programmed death mechanisms such as autophagy, pyroptosis, and ferroptosis[7]. Among them, ferroptosis is a new type of programmed cell death, and a programmed cell death pathway that relies on irons and reactive oxygen species (ROS) was reported in Cell in 2012 and named ferroptosis[8, 9]. The development of ferroptosis is closely related to many biological processes, mainly intracellular glutathione (GSH) depletion, reduced glutathione peroxidase 4 (GPX4) activity, failure of the GPX4-catalyzed reduction reaction to metabolize lipid peroxides, and oxidation of lipids by Fe^2+^ in a Fenton-like manner, generating large amounts of ROS and promoting ferroptosis[10]. As research progresses, more and more scholars are finding that iron death plays a vital role in the onset and progression of ischaemic stroke, possibly through the regulation of iron metabolism, GSH/GPX4 and lipid peroxidation, and other related pathways, which compromise the structural and functional integrity of brain tissue and lead to neuronal death and dysfunction[11-13].In addition to ischemic stroke, ferroptosis also plays an essential role in developing of many diseases, such as cancer and heart disease[14]. Therefore, genetic markers associated with ferroptosis can be used as biomarkers for diagnosing, prognosis, or treating many cancers[15]. However, the role of ferroptosis-related genes in diagnosing and treating ischemic stroke has yet to be fully elucidated.

In this study, IS microarray datasets and ferroptosis genes were obtained from the GEO and FerrDb databases. Differential gene analysis and ferroptosis-related genes(FRGs) were performed by bioinformatics analysis. Subsequently, a machine learning strategy was used to screen characteristic FRGs for IS. These FRGs were then subjected to functional enrichment, diagnostic efficacy assessment, immune cell infiltration analysis, drug target prediction, and ceRNA construction.

### Materials and methods Acquiring data

In the GEO database https://www.ncbi.nlm.nih.gov/gds/[16], search for “ischemic stroke” and select “Homo sapiens” as the species. In the search results, we selected a dataset containing blood samples from ischemic stroke patients and healthy controls, with >10 samples from experimental and control groups. GSE22255[17] was obtained as the training dataset on platform GPL570, including 20 blood samples from ischemic stroke patients and 20 control samples. GSE16561[18] was obtained as the IS biological target validation dataset on platform GPL6883, including 39 blood samples from ischemic stroke patients and 24 control samples. In addition, we obtained 380 FRGs from FerrDb[19] (http://www.zhounan.org/ferrdb).

### Identification of DE-FRGs

IS genes from the GSE22255 were intersected with the 380 FRGs to obtain IS-associated FRGs using R software. Differential expression analysis of these FRGs was performed according to the splitting criteria: P<0.05 and |log FC| >1. These differentially expressed ferroptosis-related genes (DE-FRGs) were visualized by drawing expression heatmap and correlation heatmap using R software.

### Functional enrichment analysis

We explored the biological pathways and functions of up-and down-regulated DE-FRGs using the R software package “clusterProfiler”[20]. First, the Kyoto Encyclopedia of Genes and Genomes (KEGG) assessed the biological pathways associated with differentially expressed genes. Then, three types of biological processes, such as Biological Process (BP), Molecular Function (MF), and Cellular Component (CC), were further investigated based on Gene Ontology (GO) functional analysis.

### Screen characteristic FRGs for IS

We used the least absolute shrinkage and selection operator (LASSO) algorithm based on the R software “glmnet” package and the support vector machine - recursive feature elimination (SVM-RFE) based on the “e1071” package to further screen for characteristic FRGs biomarkers associated with IS[21, 22]. The biomarkers obtained by both methods were intersected to get the characteristic FRGs and visualized by R software.

### Characteristic FRGs validation and diagnostic value assessment

We evaluated the diagnostic accuracy of characteristic FRGs by plotting receiver operating characteristic (ROC) curves on the dataset GSE22255 and GSE16561. The area under curve(AUC) was calculated, and the AUC was taken to be in the range of 0-1, with a larger AUC indicating better predictive performance. Nextly, to facilitate the derivation of the probability of disease diagnosis by the characteristic FRGs, nomogram will be drawn by the R software “rms” package. Nomogram is a quantitative analysis chart representing the functional relationship between multiple variables with a cluster of disjoint line segments in plane coordinates. Multiple line segments are drawn according to a specific proportion, and the risk probability of an individual can be conveniently calculated by making Nomogram.

### Gene set enrichment analysis

Characteristic FRGs biomarkers were classified into high and low-expression groups based on the median expression of all samples of essential gene biomarkers. Using R software, c2.cp.kegg.symbols were selected as the reference gene set for single-gene gene set enrichment analysis (GSEA) analysis. Positive gene sets were screened at a threshold of P<0.05 and a false discovery rate (FDR) <0.25.

### Immune infiltration analysis

The gene expression data of 22 immune cells were downloaded from CIBERSOFT (https://cibersort.stanford.edu/) using R software[23]. The percentage of 22 immune cells in the two groups was calculated using the “e1071” package to plot the abundance of immune infiltration. The correlation heat map and violin plot were plotted using the “corrplot” package and “vioplot” package, respectively, to analyze the correlation of immune cell infiltration distribution and its difference, with P < 0.05 representing a significant difference between the two groups.

### Predicting potential therapeutic drug for IS

Drug-Gene Interaction Database (DGIdb, http://dgidb.org/) is a database that integrates information on drug-gene interactions and gene druggability from organized and presented papers, databases, and web resources. The characteristic FRGs were uploaded to the DGIdb database for finding drug-gene action relationships. Visualization of drug-gene regulatory networks is mapped by Cytoscape software.

### Construction of ceRNA network

The miRNA target databases miRDB, TargetScan, and miRanda were used to predict the miRNAs bound by characteristic FRGs. The threshold value set was that the predicted target gene-miRNA relationship pairs appeared in all three databases, which means that the gene-regulated miRNAs were considered to have a target relationship. The spongeScan database predicted the miRNA-binding IncRNAs. Finally, the above mRNA-miRNA-IncRNA regulatory relationships were used for ceRNA network construction using Cytoscape software.

## Results

### Identification of DE-FRGs

The levels of expression of 236 FRGs were compared in the GSE22255 cohort (20 normal and 20 IS samples). Among them, we identified 22 DE-FRGs, of which 12 (DUOX2, TP53, LPCAT3, ACO1, ELAVL1, HILPDA, MIOX, PEX12, MDM2, DLD, SLC7A11, and METTL14) were downregulated, while 10 (FTH1, KRAS, EGFR, ATF3, MMD, MAP3K14, DDR2, TRIM46, DPEP1, and COX4I2) were upregulated. Based on DE-FRGs, we plotted the gene expression and correlation heatmap between the control and experimental groups(Fig 1). The results showed that the expression of 22 DE-FRGs was statistically different in both groups, and multiple gene pairs were positively and negatively regulated between the two groups.

**Fig 1.**
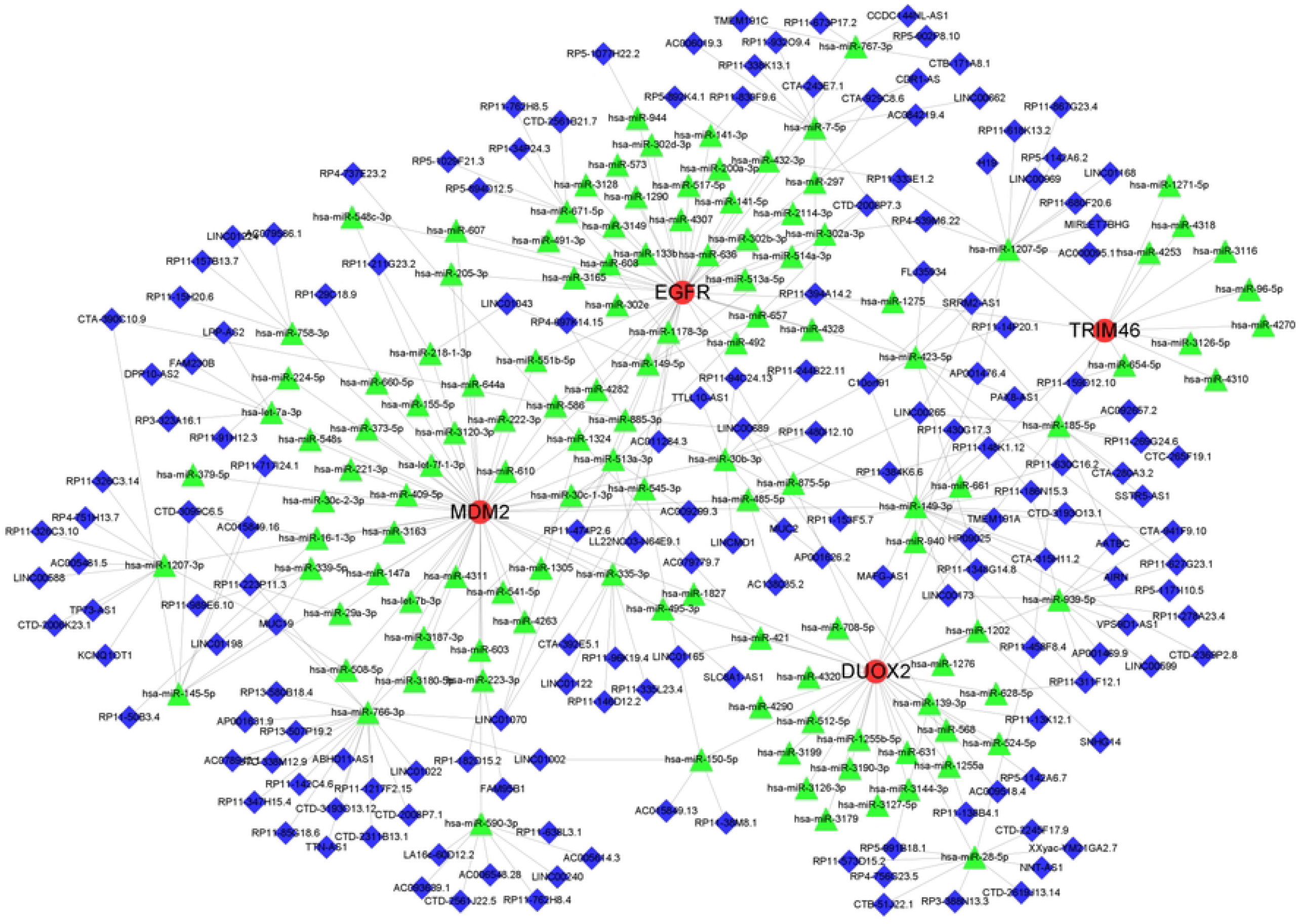
Analysis of DE-FRGs expression levels in IS. (**a**) Heat map of the 22 DE-FRGs in GSE22255 dataset. (**b**) The correlation of these genes.

### Analysis of enrichment function

We conducted GO enrichment and KEGG pathway analyses to understand better the biological activities and pathways linked DE-FRGs. GO enrichment indicated that the most enriched biological processes, cellular components, and molecular function were “cellular modified amino acid metabolic process,” “apical part of cell,” and “oxidoreductase activity, acting on NAD (P)H” (Fig 2A, B). KEGG pathway analysis was mainly enriched in signaling pathways such as the “Ferroptosis,” “MAPK signaling pathway,” and “PI3K−Akt signaling pathway” (Fig 2C, D).

**Fig 2.**
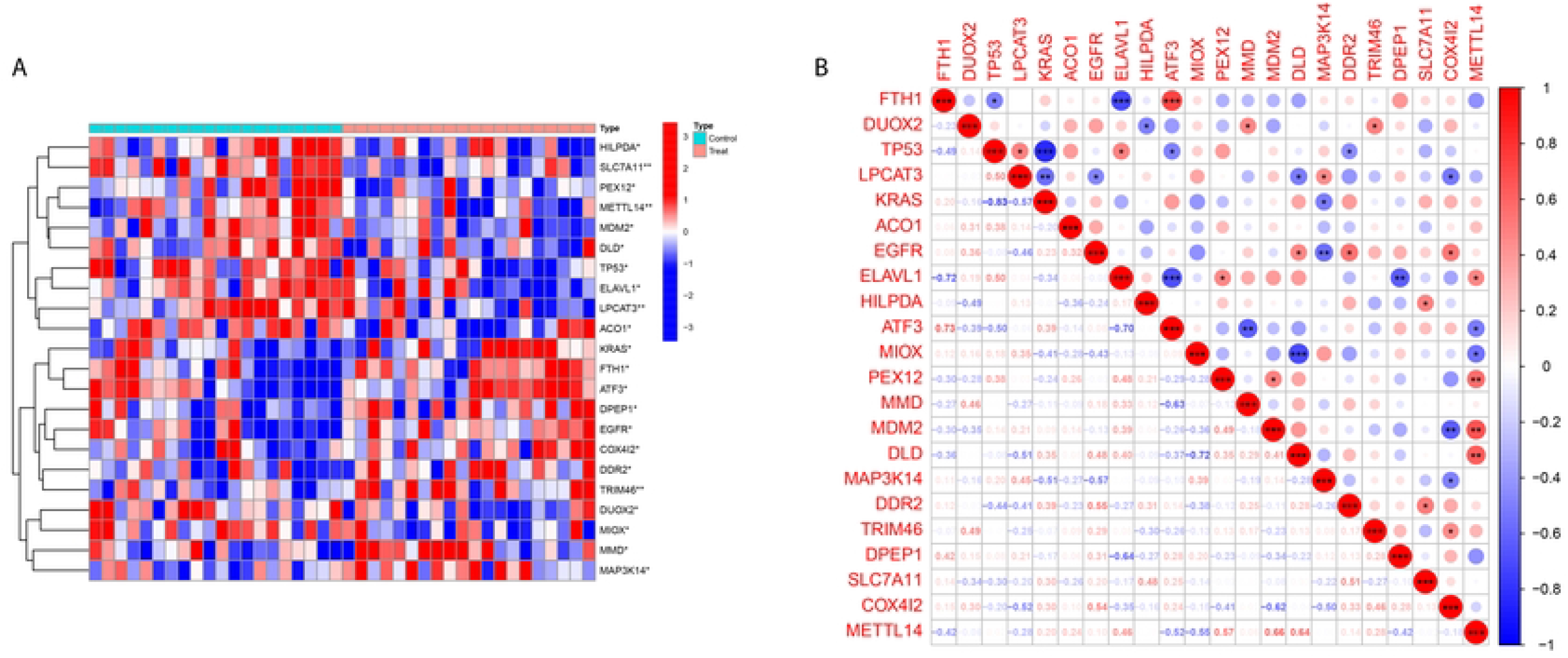
Enrichment analysis of DE-FRGs in IS. (**a) (c**) Column diagrams of GO and KEGG enrichment analysis. (**b) (d**) Circle diagrams of GO and KEGG enrichment analysis.

### Selection of characteristic FRGs

The DE-FRGs were further screened using LASSO regression and SVM-RFE algorithms, respectively. The LASSO regression model was constructed and cross-validated. The minimum error value corresponds to 12 feature *DE-FRGs*, and the SVM-RFE algorithm selected nine feature *DE-FRGs* by 10-fold cross-validation(Fig 3A-D). The feature genes obtained from both methods were intersected to finally get five characteristic FRG*s*, DUOX2, EGFR, MDM2, MAP3K14, and TRIM46, shown in Fig 3F.

**Fig 3.**
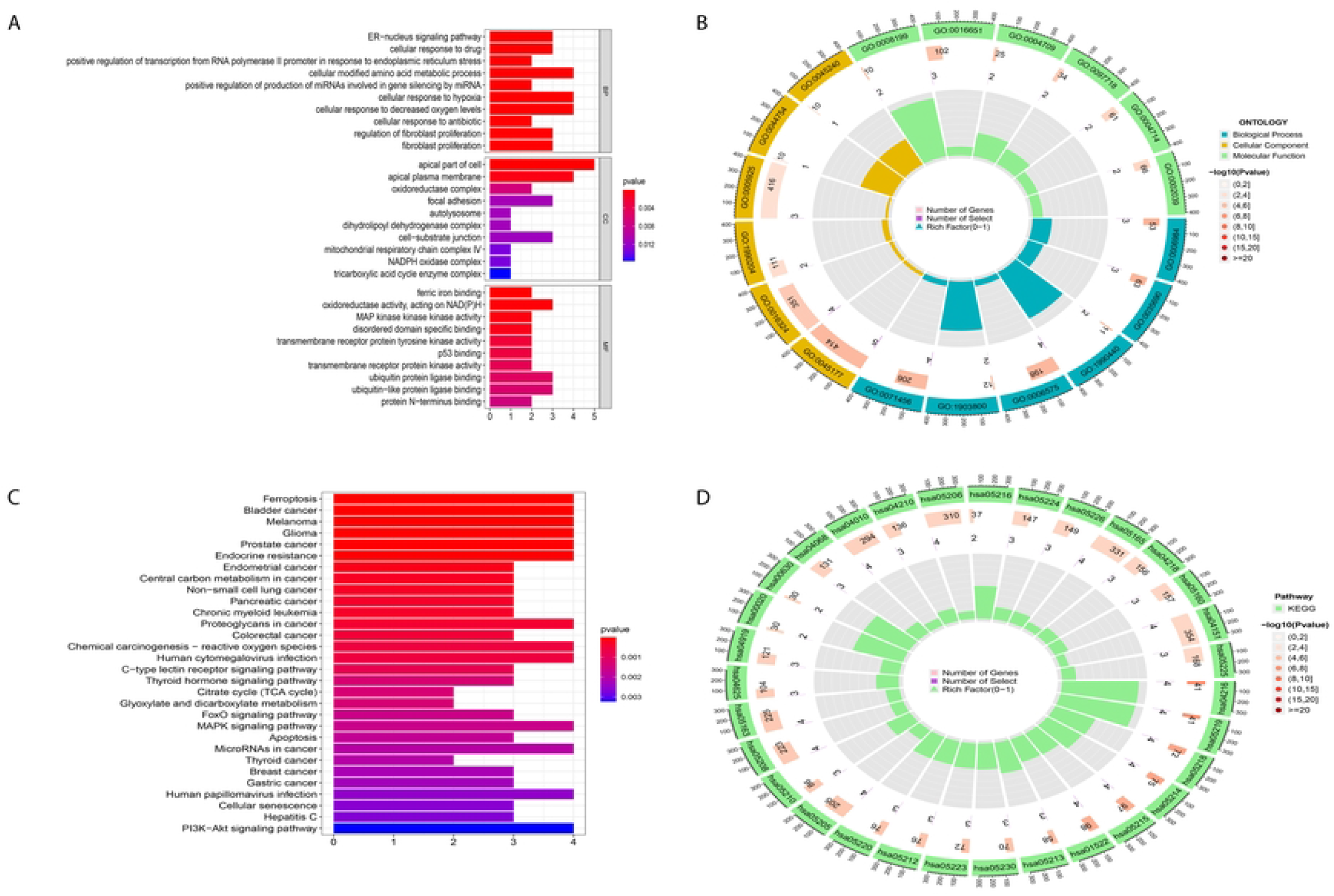
Screening for characteristic FRGs using machine learning algorithms. (**a) (b**) LASSO logistic regression algorithm to screen diagnostic markers; different colors represent different marker. (**d) (c**) Based on support vector machine-recursive feature elimination (SVM-RFE) algorithm to screen biomarkers. (**e**) Venn diagram of the LASSO and SVM-RFE models

### Validation of characteristic FRGs

To further validate the diagnostic value of the characteristic FRGs. The ROC curves showed that the AUC values of characteristic FRGs were higher than 0.7 in the GSE22255 dataset (Fig 4A). The same results were obtained in the validation dataset GSE16561, indicating that these FRGs have a high predictive effect in distinguishing IS patients from healthy controls (Fig 4B).

**Fig 4.**
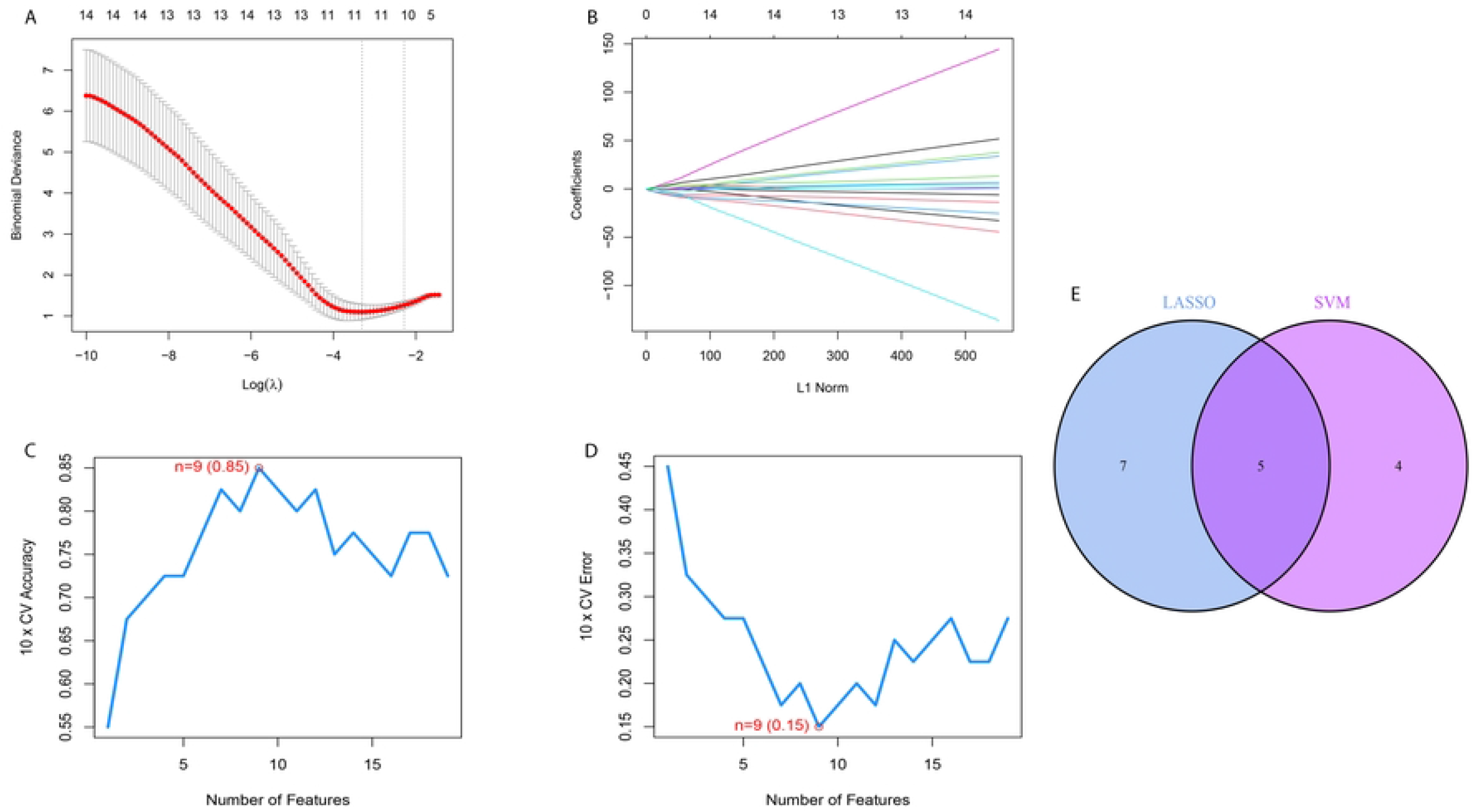
ROC analysis of characteristic FRGs. (**a**) on the GSE22255 dataset. (**b**) on the GSE16561 dataset.

### Construction and evaluation of nomogram model

Based on five characteristic FRGs expression matrices, logistic regression constructed prediction models and visualized them by drawing nomogram plots(Fig 5A). The ROC curve results showed that the C-index of the prediction model was 0.963(Fig 5B), and the constructed nomogram model had the highest performance in diagnosing IS patients compared with other single-characteristic FRG. The model was validated by the Bootstrap method with 1000 repeated samples. The results of the calibration curve analysis showed that the model predicted the incidence of the ischaemic stroke to be in general agreement with the actual incidence(Fig 5C).

**Fig 5.**
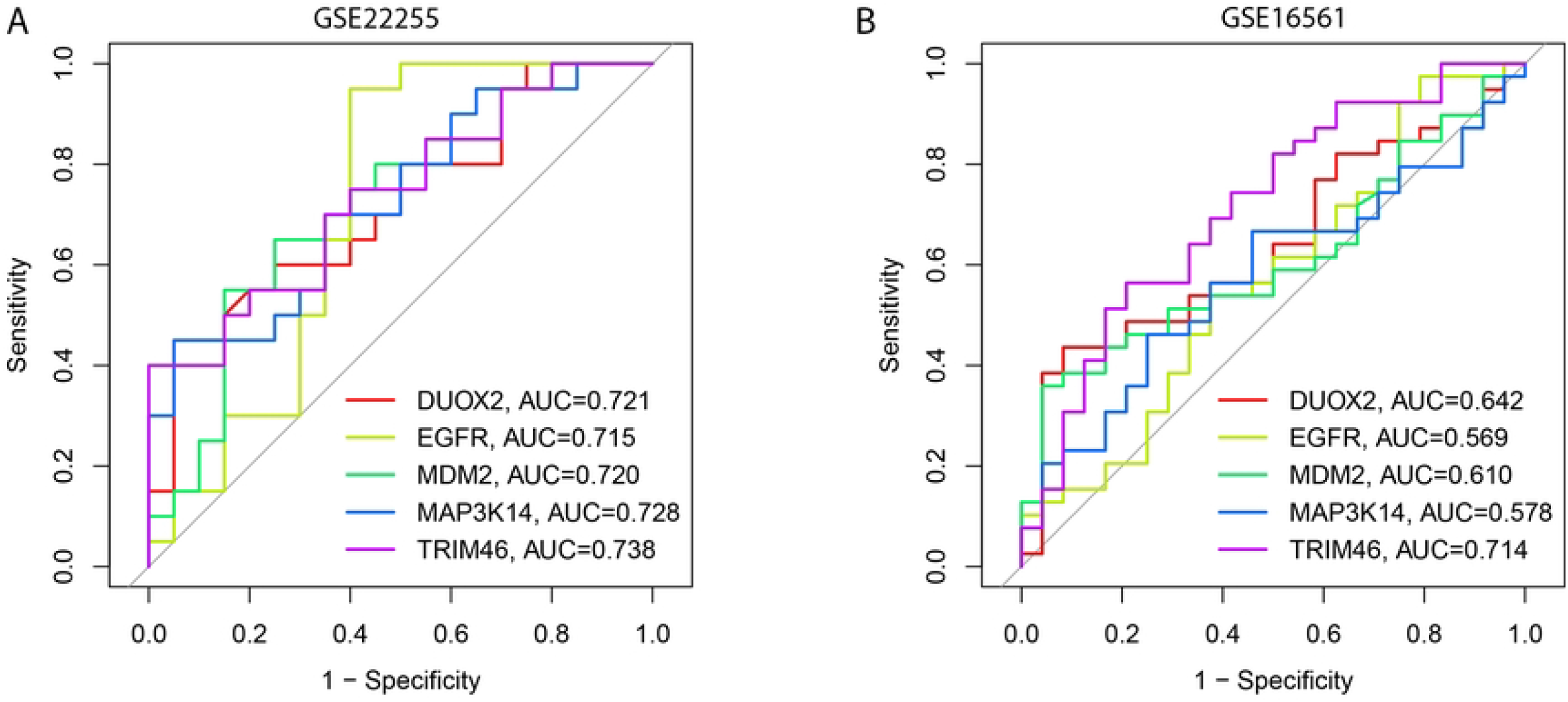
Construction and validation of a nomogram model for IS diagnosis. (**a**) The nomogram of characteristic FRGs to predict the occurrence of IS. (**b**) The ROC curve of the diagnostic efficacy verification. (**c**) The calibration curve to assess the predictive power of the nomogram model.

### GSEA for characteristic FRGs

To further explore the biological events associated with characteristic FRGs in IS, we performed a single-gene GSEA-KEGG analysis. Figure 6A-G illustrates the top 6 pathways enriched for each characteristic FRG. After a comprehensive analysis, we found that these genes are mainly enriched in the “jak stat signaling pathway,” “nod-like receptor-signaling-pathway,” “chemokine-signaling-pathway,” and “toll like receptor signaling pathway.” Also found to be associated with “graft versus host disease,” “type I diabetes mellitus,” and “systemic lupus erythematosus.”

**Fig 6.**
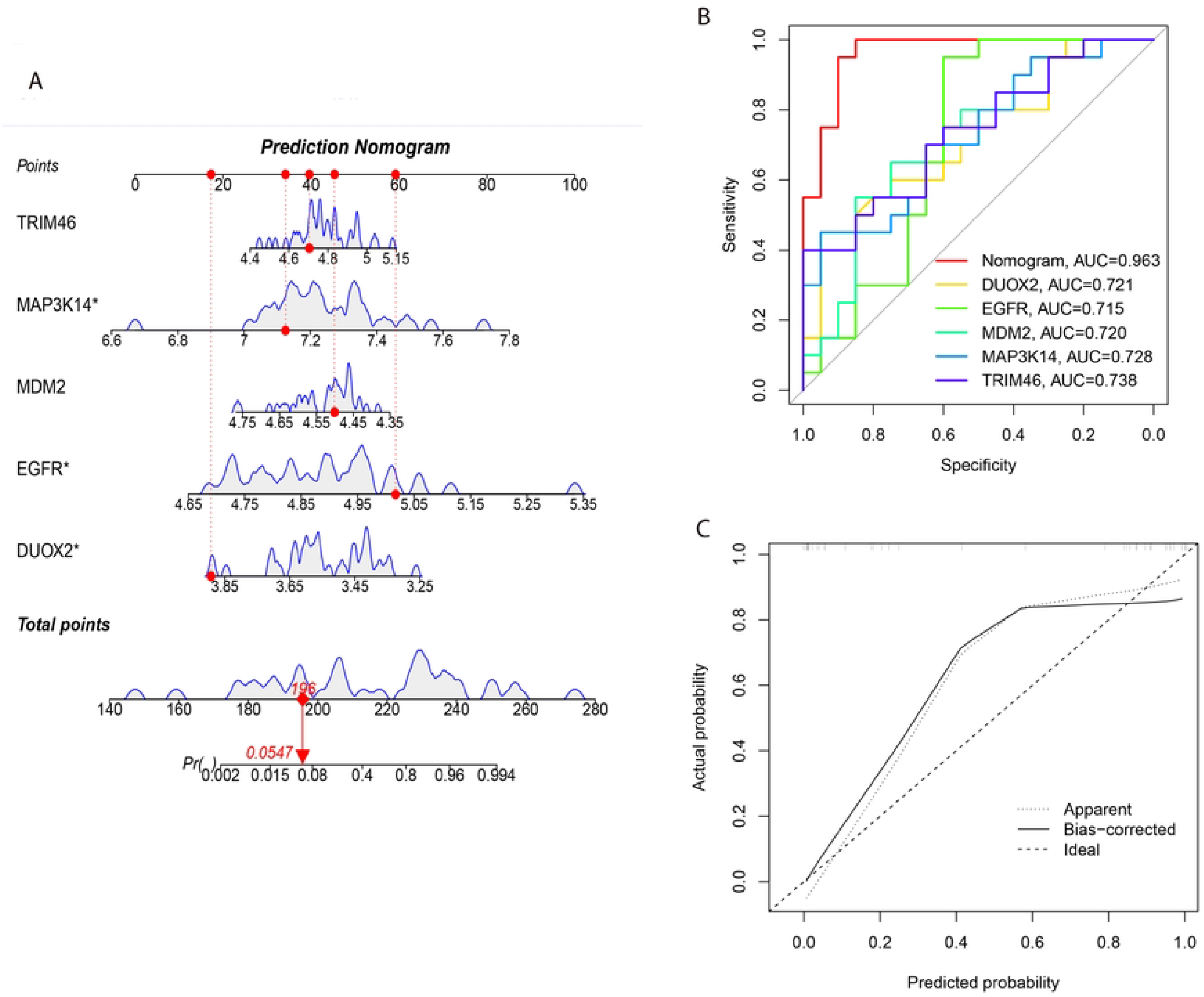
Single- gene GSEA- KEGG pathway analysis. (**a**) DUOX2. (**b**) TRIM46. (**c**) MDM2. (**d**) MAP3L14. (**e**) EGFR.

### Immune infiltrate analysis

A matrix of peripheral blood immune cell content from 20 healthy controls and 20 IS patients was constructed, showing the differences in the infiltration distribution of 22 immune cell types in different samples, as shown in Figure 7A. Immune cell correlation heatmap results show the most significant positive correlation was found by immune cell correlation analysis between “Macrophages M1” and “Dendritic cells resting” (r=0.82); the most significant negative correlation between “Mast cells activated” and “Monocytes” (r=-0.63)(Fig 7B). Violin plots were drawn to analyze the differences between the 22 immune cell infiltrations in the two groups, and only the difference in “Mast cells resting” infiltration was statistically significant. “Mast cells resting” was downregulated in peripheral blood mononuclear cells of IS patients compared to the healthy group(Fig 7C). In addition, spearman correlation analysis of 5 characteristic FRGs with 22 immune cells showed that TRIM46 had a positive correlation with” Neutrophils “and “T cells CD4 memory activated” (P<0.05), respectively, and MDM2 had a negative correlation with “Dendritic cells activated” (P<0.05); DUOX2 positive correlation (P<0.05) with “B cells memory,” “Monocytes” (Fig 7D). This evidence suggests that changes in the immune microenvironment of IS patients may be related to TRIM46, MDM2, and DUOX2.

**Fig 7.**
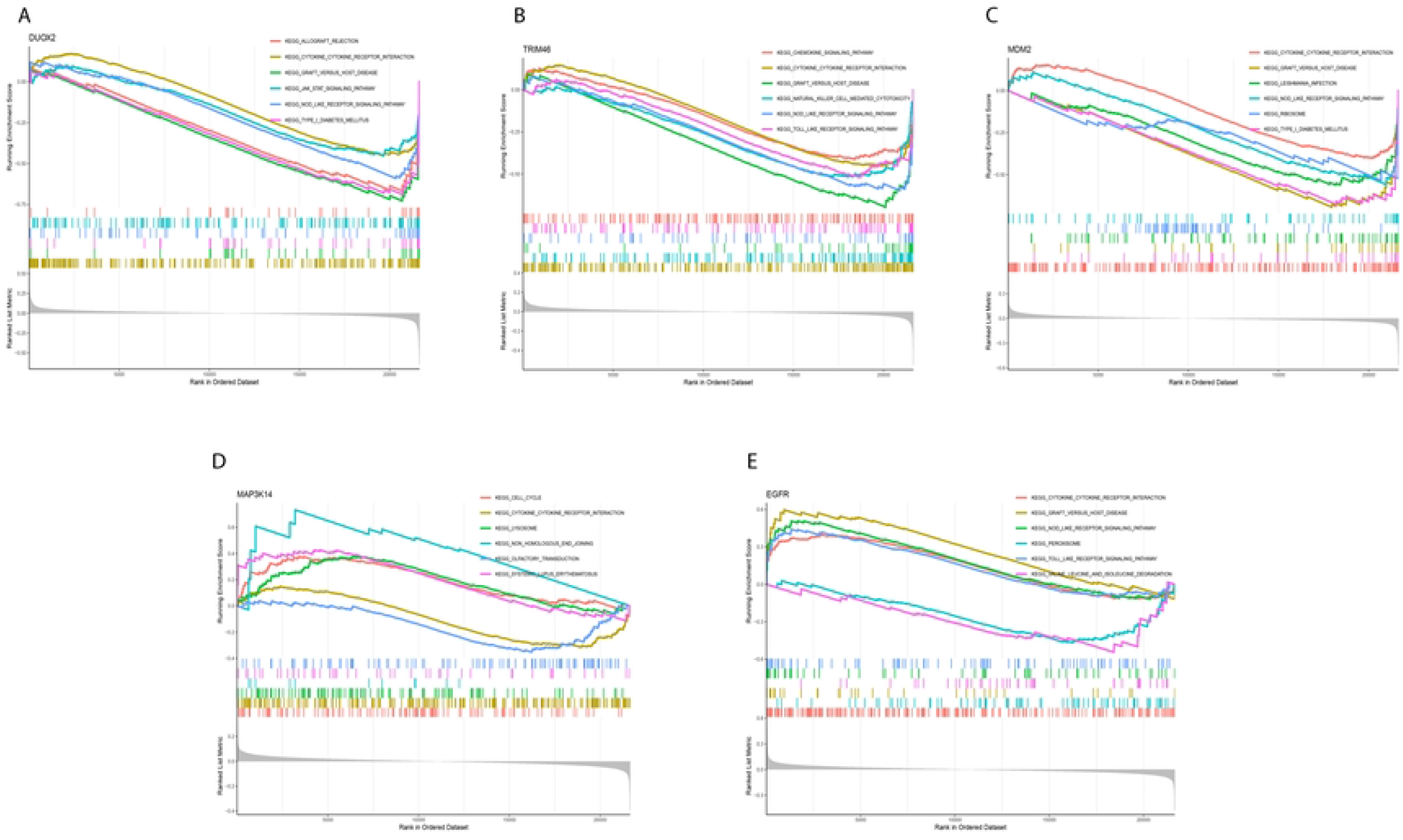
Results of immune infiltration by CIBERSORT. (**a**) Bar-plot showed the composition of 22 kinds of immune cells in GSE22255. (**b**) Correlation heatmap of 22 immune cell subpopulations. Red and blue represent positive and negative correlation, respectively. The rectangle with a deeper color has a stronger correlation index.(**c**) Violin diagram presented the difference of immune infiltration of 22 kinds of immune cells. (**d**) Correlation heatmap of immune cell and characteristic FRGs.

### Prediction of characteristic FRGs-targeting drugs

We used the DGIdb database to reveal drugs that may target the characteristic FRGs. We predicted 66 drugs targeting key FRGs, of which 50 were for EGFR and 16 for MDM2. However, we did not forerun drugs targeting DUOX2, MAP3K14, and TRIM46. Among them, DACOMITINIB, AZD-4769, AFATINIB, SAPITINIB, NERATINIB, POZIOTINIB, OSIMERTINIB, TAK-285, AC-480, VANDETANIB, CHEMBL3397300, VARLITINIB, MP-412, AZD-3759, MAB-425, ROCILETINIB, CUDC-101, LIFIRAFENIB, and CANERTINIB are inhibitors of EGFR, while RO-5045337 is an inhibitor of MDM2. The results were visualized using Cytoscape software to create a network diagram(Fig 8).

**Fig 8.**
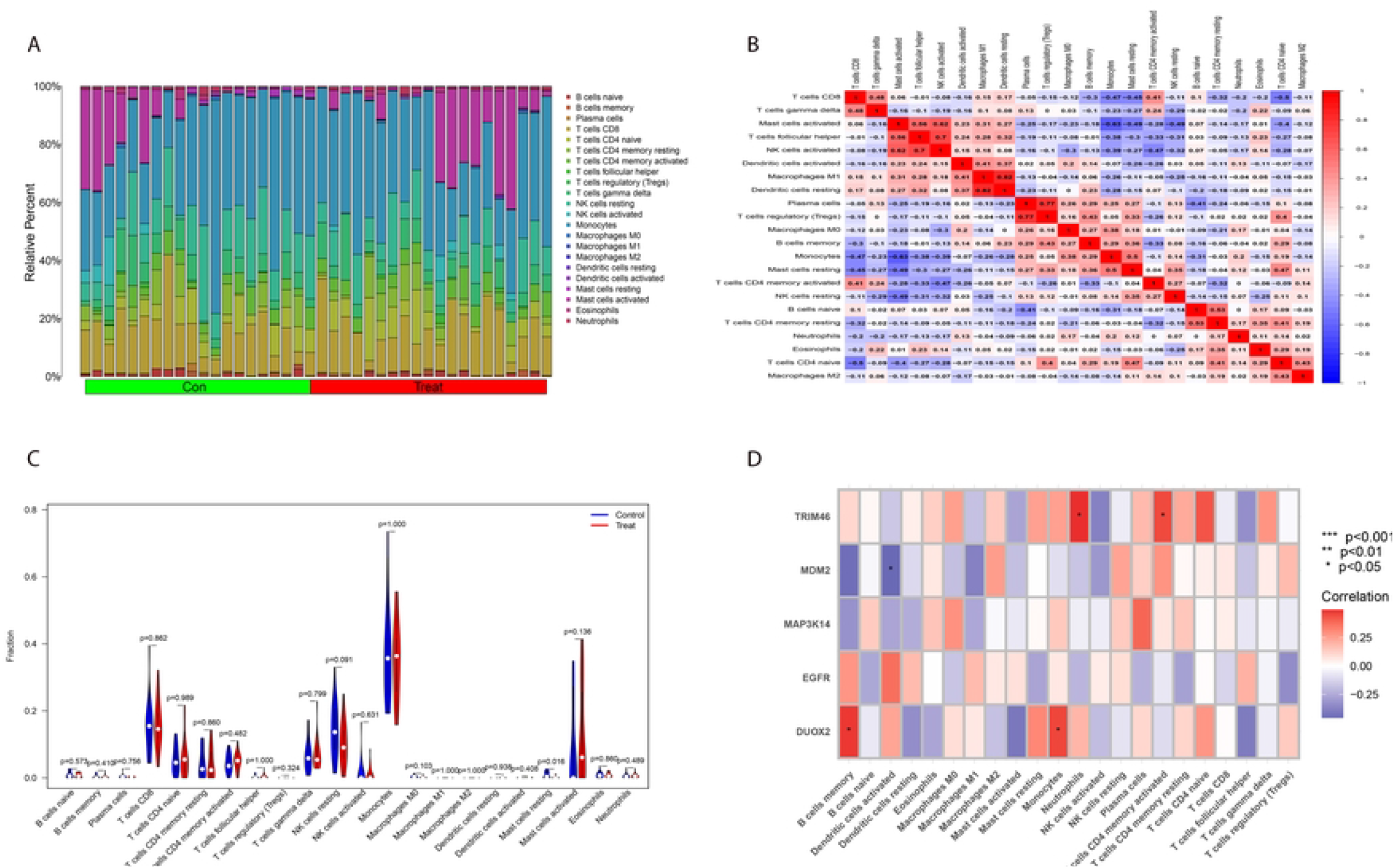
Prediction of characteristic FRGs-targeted drug. The drug-gene network of EGFR and MDM2.

### Construction of ceRNA networks based on characteristic FRGs

Based on the characteristic FRGs, 139 mRNA-miRNA pairs, and 221 miRNA-lncRNA pairs were predicted. In terms of these results, a ceRNA network of mRNA-miRNA-lncRNA was established by Cytoscape software, involving 295 nodes (4 mRNA, 124 miRNAs, and 167 lncRNAs) and 360 edges (Fig 9).

**Fig 9.**
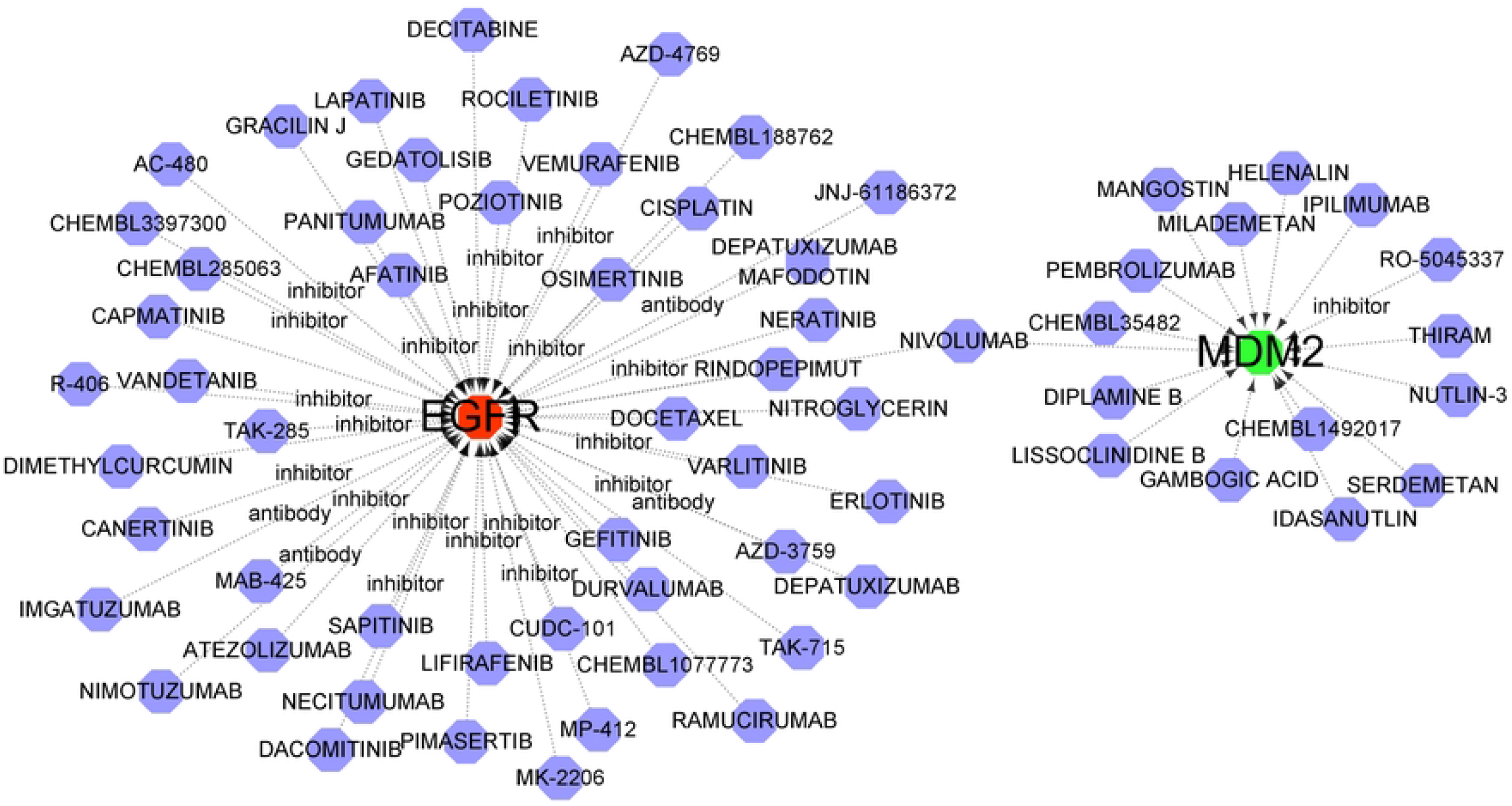
A ceRNA network based on characteristic FRGs. The red circles represent characteristic FRGs, the green triangles represent miRNAs and the blue diamonds represent lncRNAs.

## Discussion

We identified 22 DE-FRGs in the peripheral blood of individuals with ischemic stroke. Subsequently, functional enrichment analysis of these DE-FRGs was performed, and KEGG results showed that the enrichment was mainly in the MAPK signaling pathway and PI3K/Akt signaling pathway. Studies have shown that the MAPK signaling pathway is closely related to inflammation-mediated ischemic injury[24]. The MAPK signaling pathway inhibition can inhibit neuronal apoptosis and inflammatory response, protect damaged brain tissue, and improve nerve injury[25]. Many studies have also shown that PI3K/Akt signaling pathway is essential in the pathogenesis of ischemic stroke and plays a vital role in the anti-apoptosis of nerve cells, oxidative stress, inflammation, inflammation, and inflammation autophagy[26-28].

This study screened five characteristic genes related to ferroptosis, including DUOX2, EGFR, MDM2, MAP3K14, and TRIM46. The AUC value represented by the area under the ROC curve of the five genes was all greater than 0.5. Meanwhile, the constructed column graph also had high diagnostic values, indicating that these five genes had excellent accuracy and specificity in distinguishing IS disease samples from standard samples. Dual oxidase 2(DUOX2) belongs to the nicotinamide adenine dinucleotide phosphate (NADPH) family of oxidases, which consists of seven members: Nox1, Nox2, Nox3, Nox4, Nox5, Duox1, and Duox2. The Nox family of NADPH oxidases has been shown to play an important role in cerebral ischemia-reperfusion injury by causing oxidative stress, nitrative stress, blood-cerebrospinal fluid destruction, inflammatory responses, and apoptosis. DUOX2, located on human chromosome 15, was initially identified in thyroid tissue and is associated with thyroid dysfunction and congenital hypothyroidism. It has been shown that DUOX2 plays different roles in the development of various malignancies, and GSEA analysis has also revealed that DUOX2 may be involved in tumor-related signaling pathways, such as the JAK-STAT signaling pathway. However, there are no reports on the role of DUOX2 in the pathogenesis of IS, and further studies are needed. Epidermal growth factor receptor (EGFR) is a typical tyrosine kinase receptor composed of an intracellular ligand-binding region, an extracellular region, and a transmembrane area[29]. It is widely involved in various pathophysiological processes, such as cell differentiation, proliferation, migration, invasion, and damage repair. EGFR is not detected in astrocytes in the average adult central nervous system. However, EGFR reappears in reactive astrocytes after injuries such as ischemia and trauma[30]. Studies have found that EGFR inhibitors can inhibit EGFR phosphorylation of alien glial cells in vivo, reduce the proliferation of astrocytes after ischemia, reduce neuronal apoptosis, reduce infarct volume, and promote behavioral recovery by establishing a rat model of cerebral ischemia in vivo[31]. Studies have demonstrated that activation of EGFR can increase the proliferation and survival of neural precursor cells and reduce ischemic brain injury[32]. TRIM46 (tripartite motif - containing 46, TRIM46) is to TRIM the 46 members of the family on chromosome 1 q21, composed of 759 amino acids[33]. TRIM46 is widely expressed in various tissues, especially in the brain, such as in the first segment of neuronal axons[34]. Recent studies have found that TRIM46 is also associated with ROS content in human retinal capillary endothelial cells, suggesting that TRIM46 may regulate cell ferroptosis[35]. However, relevant research has yet to be found on TRIM46 regarding IS, which needs to be further developed.

Energy expenditure and hypoxia after IS can lead to nerve cell necrosis, which activates resident glial cells and promotes peripheral immune cell invasion into the ischemic brain[36]. Recent studies have shown that immune cell-mediated inflammatory response plays an important role in the pathogenesis of IS[37-39]. The injury-related molecular model molecules released by damaged neurons after the occurrence of IS can trigger the local immune response, leading to the activation of glial cells and the recruitment of peripheral white blood cells to the damaged brain area, and then the secretion of a variety of pro-inflammatory cytokines, chemokines, and matrix metalloproteinases, leading to the damage of the blood-brain barrier, brain edema, hemorrhage transformation, and neuron necrosis[40]. After cerebral ischemia, mast cells release vasoactive substances and inflammatory mediators (such as histamine, protease, TNF-α, *Etc*.), leading to an imbalance of inflammatory state[41]. Mast cells also regulate blood - brain barrier permeability, and T cells and myeloid cells infiltrate the central nervous system and have phagocytosis and antigen presentation functions [42]. This study used the CIBERSORT algorithm to assess the type of immune cell infiltration in IS and standard peripheral blood samples. The results showed that resting mast cell infiltration decreased in IS samples compared with healthy people, consistent with previous studies. At the same time, we analyzed the correlation between characteristic FRGs and immune cells. However, how these FRGs regulate immune cells is still unclear, which needs continuous attention and exploration in subsequent clinical trials.

To further explore the potential therapeutic perspective of these characteristic FRGs on IS, we used the DGIdb database to search for medicinal drugs or molecular chemicals that can interfere with IS. Most of these drugs mentioned are associated with tumor diseases and have not been reported in the application of IS. Further studies are needed to evaluate the ability of the clinical application of IS. Our study also focused on non-coding RNAs associated with these critical genes. Previous studies have shown that non-coding RNAs play an essential regulatory role in IS through various pathways, such as apoptosis, oxidative stress, activation of microglia, and regeneration of vascular endothelial cells[43]. As an important regulatory factor of IS, miRNA is involved in the occurrence and regulation of ischemic nerve tissue inflammation and has potential application value for the neuroprotection and neovascularization of IS[24]. Ischemic stroke can cause significant changes in the lncRNA expression profile. Thousands of abnormally expressed lncRNAs have been found in patients with ischemic stroke and animal models [44]. LncRNA affects the progression and outcome of ischemic stroke through calcium overload, inflammatory response, neurovascular injury, neuronal necrosis, neuroprotection and repair, and synaptic function destruction, and participates in the molecular process of ischemic cascade[45]. We constructed a ceRNA network with 295 nodes (4 mRNAs, 124 miRNAs, 167 lncRNAs) and 360 edges, providing a basis for further study of IS.

## Conclusions

In this study, five ferroptosis-related diagnostic markers of IS were obtained. We constructed the diagnostic model, the role of related immune landscape and immune cells in IS, and explored the corresponding mechanism. The drug network and ceRNA related to IS treatment were constructed. The disadvantage is that we need to verify it through clinical samples or basic experiments. It provides a new reference for this disease’s diagnosis, mechanism research, and treatment.

## Data availability statement

The datasets used and analyzed during the current study are available from the First author on reasonable request.

## Authors’ Contributions

Conceptualization: Xin Feng.

Data curation: Xin Feng, Chunhua Huang.

Formal analysis: Chunhua Huang.

Methodology: Lin Ye.

Software: Chunhua Huang.

Supervision: Zhongbo Xu.

Validation: Lin Ye.

Writing – original draft: Lin Ye, Xin Feng.

Writing – review & editing: Zhongbo Xu.

## Acknowledgments

The work involved in this manuscript was supported by the Science and Technology Research Project of Jiangxi Provincial Education Department (No. GJJ2200939).

